# Temporal persistence of admixed phenotypes in a damselfly hybrid zone

**DOI:** 10.64898/2026.05.29.728807

**Authors:** María Isabel Velásquez-Vélez, Rosa Ana Sánchez-Guillén, Laura Pulido-Rios, Anwar Medina-Villarreal, Adolfo Cordero-Rivera, Emilio Realpe, Clara Inés Saldamando-Benjumea

## Abstract

In odonates, hybridization has been documented in several species pairs, yet the long-term persistence of hybrid phenotypes remains poorly understood, particularly in Neotropical systems. Here, we investigate a putative hybrid zone between *Ischnura capreolus* and *Ischnura cyane,* which overlap along an altitudinal gradient in the Colombian Andes. We combined temporal sampling across sympatric and allopatric localities with multilocus genetic analyses and geometric morphometrics of wings and male caudal appendages to evaluate patterns of admixture and phenotypic intermediacy. Morphometric analyses revealed pronounced differentiation between *I. capreolus* and *I. cyane*, especially in allopatric populations, whereas individuals from sympatric localities displayed increased morphological overlap and intermediate phenotypes. Genetic analyses based on nuclear and mitochondrial genes identified two main genetic clusters corresponding to the parental species, with evidence of admixture and shared haplotypes in sympatric localities. Patterns of differentiation varied among loci, with mitochondrial markers showing broader haplotype sharing than nuclear loci. Across sympatric localities, morphologically intermediate individuals remained consistently present through time, particularly in Anolaima, where they represented the dominant phenotype across sampling periods. These results support concordant morphological and genetic evidence consistent with persistent hybridization and introgression between *I. capreolus* and *I. cyane* in their zone of contact and suggest that incomplete reproductive isolation contributes to the long-term maintenance of admixed phenotypes in this hybrid zone.

## INTRODUCTION

In a comprehensive review of hybrid zones, Harrison (1990) proposed that hybridization during diversification rarely occurs in animals. Subsequent work has shown that hybridization is more common than previously thought (Taylor & Larson, 2019). Hybrid zones have been defined as “windows on the evolutionary process” (Harrison, 1990) and as “natural laboratories” (Barton & Hewitt, 1985) for studying the evolution of reproductive barriers (e.g., Sánchez-Guillén et al., 2012), the role of selection in maintaining species differences (e.g., Harrison & Larson, 2014), and how phenotypic traits diverge between hybridizing populations (e.g., Abbott et al., 2013).

Across taxa, hybridization can have diverse evolutionary consequences. The importance of hybridization in plant evolution is well-documented, with 30-70% of flowering plants showing evidence of past hybridization events (Soltis & Soltis, 2009). In animals, hybridization appears to be less frequent (around 10%), although estimates remain uncertain and vary among taxa (Barton, 2013). Historically, animal hybrid zones have been studied primarily to understand reproductive isolation and species boundaries. Traditional approaches have identified hybrids based on their morphological intermediacy relative to the parental species. However, some hybrid traits resemble one parental species or even exceed parental ranges through transgressive segregation (Dittrich-Reed & Fitzpatrick, 2013), particularly in later-generation hybrids and introgressed individuals (Rieseberg *et al*., 2007). More recently, multilocus and genomic approaches have improved our understanding of the evolutionary consequences of hybridization and introgression across diverse taxa. Despite this extensive body of work, the evolutionary outcomes of hybridization, particularly the long-term persistence of admixed populations and hybrid phenotypes remain incompletely understood (Abbott *et al*., 2013; Peñalba *et al*., 2024).

Hybridization and introgression have been documented across multiple evolutionary scales in odonates. At the macroevolutionary level, Suvorov *et al.,* (2022) detected that the phylogenetic history of odonates includes introgressive hybridization, defined as the incorporation of alleles from one species into the gene pool of another through hybridization and backcrossing (Anderson & Hubricht, 1938). Introgression has been identified across deep phylogenetic levels, including events occurring between major odonate lineages and among distantly related families (Suvorov *et al*., 2022). At the microevolutionary level, hybridization in odonates has been associated with factors such as climate change (Sánchez-Guillén *et al*., 2013, 2014c), community composition (Lorenzo-Carballa *et al*., 2014; Sánchez-Guillén *et al*., 2016), color polymorphism (Johnson, 1975; Ordaz-Morales *et al*., 2026), reinforcement of reproductive isolation (Tynkkynen *et al*., 2008a; Arce-Valdés *et al*., 2025), and reproductive character displacement (Ballén-Guapacha *et al*., 2024; Ballén-Guapacha & Sánchez-Guillén, 2026). Their mating system also generates mechanical and tactile incompatibilities that can act as strong prezygotic barriers to hybridization, as documented in several genera, including *Ischnura* (Monetti *et al*., 2002; Sánchez-Guillén *et al*., 2012), *Argia* (Nava Bolaños *et al*., 2016), *Enallagma* (Barnard *et al*., 2017) and *Cordulegaster* (Solano *et al*., 2018). The coexistence of introgression and well-documented reproductive barriers makes odonates particularly suitable systems for investigating how hybridization persists despite incomplete reproductive isolation. This is especially evident among closely related species within the genus *Ischnura*, which comprises around 75 species distributed worldwide (Sánchez-Guillén *et al*., 2020). Hybridization has been documented in at least 6 species pairs across Nearctic and Palearctic regions (Sánchez-Guillén *et al*., 2022).

In this study, we investigated hybridization between the damselflies *Ischnura capreolus* and *Ischnura cyane* in the Colombian Andes using an integrative approach that combined wing and genitalia morphology with nuclear and mitochondrial gene sequences. These species overlap in their altitudinal ranges, from 1300 to 1800 m a.s.l., in the Colombian Andes. In sympatric areas, around 1600 m a.s.l., morphologically intermediate localities have been reported, characterized by adults and larvae with intermediate morphology (Realpe, 2010; Galindo-Ruiz *et al*., 2019). Males of *I. capreolus* are predominantly green (Vilela *et al*., 2017) whereas *I. cyane* are mainly blue (Realpe, 2010), while putative hybrids exhibit intermediate thoracic and abdominal coloration and intermediate genital morphology (Pulido-Rios, 2023). Additionally, larval development differs among taxa, with putative hybrids showing up to 12 larval instars, compared to 10-11 in *I. capreolus* and 8 in *I. cyane* (Galindo-Ruiz *et al*., 2019). These observations suggest that sympatric localities contain individuals with intermediate phenotypic characteristics relative to the parental species. These differences suggest that hybridization may contribute to increased developmental and phenotypic variation in sympatric populations, a pattern that has been associated with transgressive phenotypes in other hybrid systems (Rieseberg *et al*., 1999; Dittrich-Reed & Fitzpatrick, 2013).

To evaluate whether patterns of genetic and morphological variation are consistent with hybridization and introgression between *I. capreolus* and *I. cyane*, we combined temporal sampling across a 12-year period with multilocus genetic analyses and geometric morphometrics of wings and male caudal appendages. Specifically, we tested whether genetic and morphological data supported concordant patterns of differentiation and admixture between taxa. We predicted that: (1) parental species and putative hybrids would be distinguishable based on genetic and morphological data; (2) hybrids would exhibit intermediate characteristics relative to the parental species; and (3) that differentiation between species would be reduced in sympatric relative to allopatric populations, consistent with introgression between taxa.

## MATERIALS AND METHODS

### Morphological traits used for species discrimination

In *Ischnura,* body coloration (thoracic and abdominal patterns) is commonly used for field identification; however, these traits may overlap among closely related species and putative hybrids, particularly in contact zones (Sánchez-Guillén *et al*., 2005). In the study system, *Ischnura capreolus* and *I. cyane* differ in their typical coloration, males being predominantly green and blue, respectively; however, putative hybrids exhibit intermediate colour patterns, which can limit the reliability of colour-based identification in sympatric localities (Pulido-Rios, 2023). Male caudal appendages and female prothorax are involved in tandem formation, and have been associated with mechanical incompatibilities between species (Robertson & Paterson, 1982; McPeek *et al*., 2009; Sánchez-Guillén *et al*., 2014b; Wellenreuther & Sánchez Guillén, 2015). Wing morphology although not directly involved in mating, provides complementary information on overall morphological differentiation among taxa (Stand-Pérez et al. under rev.). Therefore, combining caudal appendages with wing morphology provides complementary information to distinguish parental species and identify morphologically intermediate individuals, capturing variation in both reproductive and non-reproductive traits.

### Field sampling sites

The three sympatric localities (El Colegio, San Francisco and Anolaima) in the Colombian Andes (Fig. 1) were sampled five times over a 12-year period (2004-2016) to estimate the relative frequency of each species and putative hybrids. Individuals were classified in the field based on diagnostic morphological traits, including body coloration, male caudal appendages and female prothorax. Differences in taxon composition among localities were assessed using chi-square tests based on observed frequencies across sampling periods.

**Figure 1.**
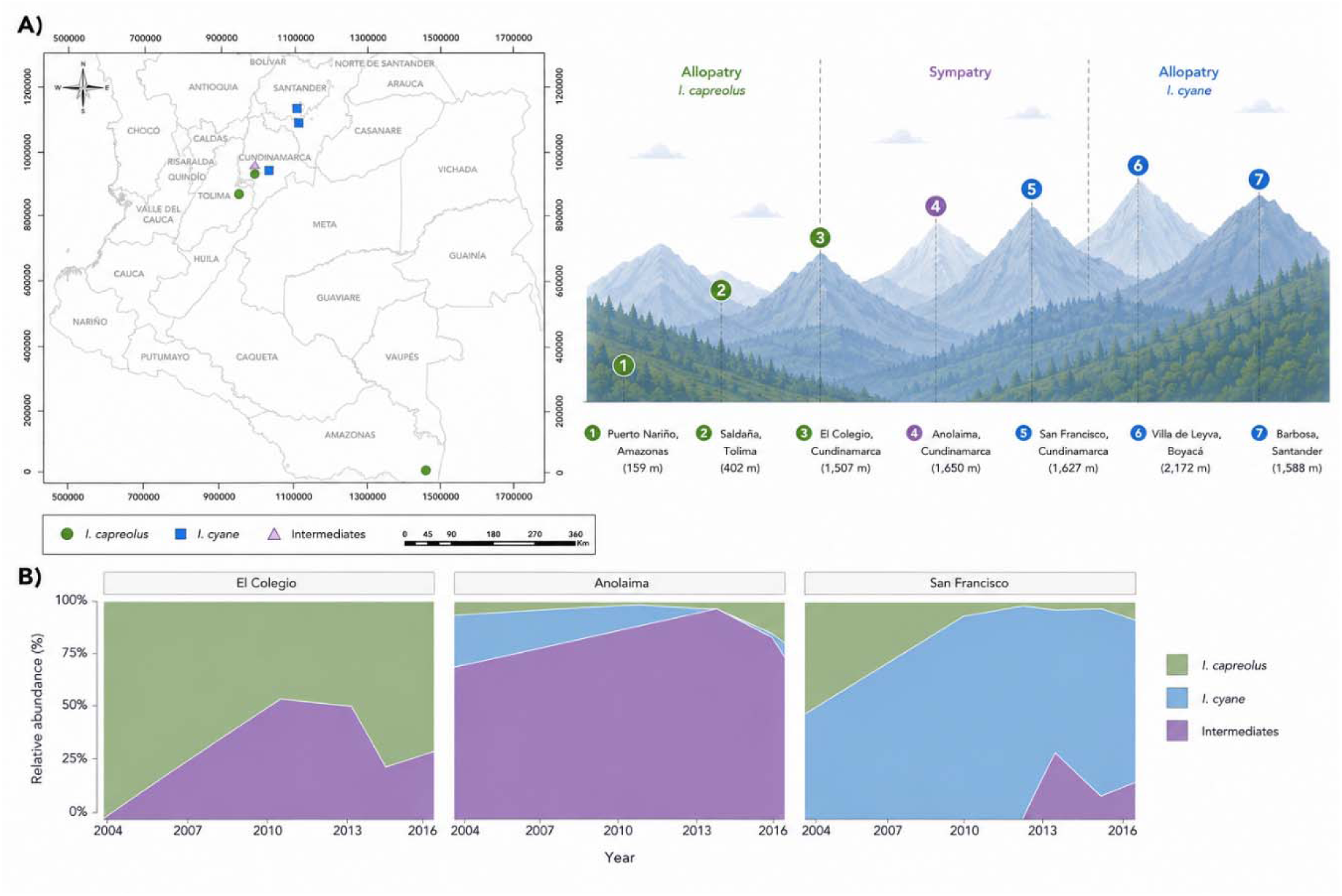
Geographic distribution and temporal persistence. **(A)** Sampling localities in Colombia, including two allopatric localities of *I. capreolus* (Puerto Nariño and Saldaña), two allopatric localities of *I. cyane* (Villa de Leyva and Barbosa), and three sympatric localities (El Colegio, San Francisco, and Anolaima) where both species and morphologically intermediate individuals occur in varying proportions. **(B)** Temporal variation in the proportion of *I. capreolus*, *I. cyane*, and morphologically intermediate individuals across the three sympatric localities from 2004 to 2016. Colors indicate taxa: *I. capreolus* (green), *I. cyane* (blue), and morphologically intermediate individuals (purple).

For the genetic and morphological analyses, adults of *I. capreolus*, *I. cyane* and morphologically intermediate individuals were sampled between 2013-2016 in seven localities of the Colombian Andes (Fig. 1): two allopatric localities of *I. capreolus* (Saldaña and Puerto Nariño), two allopatric localities of *I. cyane* (Barbosa and Villa de Leyva), and three sympatric localities: El Colegio, where most individuals were identified as *I. capreolus*, San Francisco, where most individuals were identified as *I. cyane,* and Anolaima, where most individuals were morphologically intermediate (putative hybrids). A total of 237 males and 98 females were collected and preserved in absolute ethanol (Table S1).

### DNA extraction, PCR amplification, and sequencing

A total of 150 individuals of *I. capreolus*, *I. cyane*, and putative hybrids were used for genetic analyses. Genomic DNA was extracted using the DNeasy® kit (Qiagen) according to the manufacturer’s protocol. Thoracic muscle tissue was incubated in 1.5 mL tubes containing 100 µL of TNES buffer (50 mM Tris pH 7.5, 400 mM NaCl, 20 mM EDTA, 0.5% SDS) and 1.7 µL proteinase K (10 mg/mL) following Sánchez-Guillén *et al*. (2014a).

PCR amplifications were performed in 10 µL reaction volumes containing 2 ng genomic DNA (2 µL), 5 µL 2× Ready Mix PCR Master Mix (1.5 mM MgCl), 1 µL BSA, 0.3 µL MgCl (50 mM), 1.1 µL distilled water, and 0.3 µL of each primer (10 pmol). Nuclear (28S, ITS, and PRMT) and mitochondrial (COI, 16S, ND1, and Cytb) genes were amplified using primers and annealing temperatures listed in Table S2. Differences in sample size among loci resulted from variation in amplification success (Table S1).

PCR cycling conditions consisted of an initial denaturation at 94 °C for 4 min; 35 cycles of 94 °C for 30 s, locus-specific annealing temperature for 45 s, and 72 °C for 45 s; followed by a final extension at 72 °C for 6 min. PCR products were purified using exonuclease treatment (Bioline), then incubated at 72 °C for 15 min and at 85 °C for 15 min. Amplified products were bidirectionally sequenced by Macrogen Inc. Sequences were manually edited and assembled in Geneious v5.3.6 (Biomatters) and aligned using the ClustalW algorithm implemented in Geneious.

### Genetic analyses of structure and admixture

To assess patterns of genetic differentiation and admixture between *I. capreolus*, *I. cyane* and morphologically intermediate individuals we analyzed a combination of mitochondrial (COI, 16S, ND1, and Cytb) and nuclear (28S, ITS, and PRMT) markers. This multilocus approach was used to compare patterns of variation across marker types with different modes of inheritance. Comparing mitochondrial and nuclear markers also allowed us to evaluate whether patterns of differentiation and haplotype sharing were consistent across loci. Because mitochondrial and nuclear genes differ in their inheritance and evolutionary dynamics, they may also differ in their patterns of differentiation and introgression (Harrison & Larson, 2014; Swaegers *et al*., 2022).

Genetic diversity within populations was estimated in DnaSP v5 (Librado, P., & Rozas, J. (2009) using nucleotide diversity (π), number of segregating sites (S), number of haplotypes (h), and haplotype diversity (Hd). These indices were used to characterize and compare levels of genetic variation across markers. Population genetic structure was analyzed using AMOVA and pairwise F_ST_ estimates implemented in Arlequin 3.5 to quantify genetic differentiation among populations grouped according to species identity and geographic context (allopatry vs. sympatry). Pairwise F_ST_ values were also used to determine whether sympatric populations exhibited lower genetic differentiation than allopatric populations, consistent with introgression between taxa.

Bayesian clustering analyses were performed in STRUCTURE v2.3.4 (Pritchard *et al*., 2000) to infer genetic clustering and admixture among individuals and localities. Sequence alignments were converted into genotype matrices in which each polymorphic site was coded as an independent marker. Given the limited number of loci available, this approach was retained to maximize the number of informative sites used to detect population structure and admixture (Haasl & Payseur, 2011). Because some polymorphic sites originated from linked mitochondrial loci, STRUCTURE outputs were interpreted as exploratory estimates of clustering and admixture rather than precise measures of ancestry proportions. STRUCTURE analyses were conducted for K = 1–10 using 200,000 MCMC iterations after a burn-in period of 15,000 iterations, with five independent replicates for each K value. The most likely number of genetic clusters was inferred using the Evanno method implemented in STRUCTURE HARVESTER (Earl & vonHoldt, 2012).

Haplotypes were identified in DnaSP and visualized as median-joining networks in POPART (Leigh & Bryant, 2015) to examine patterns of haplotype sharing among species and populations. These analyses were used to evaluate whether sympatric populations showed evidence of haplotype sharing between taxa.

### Morphometric analyses of morphological differentiation

To evaluate morphological differentiation and identify morphologically intermediate individuals based on shape variation, we performed geometric morphometric analyses of wings and male caudal appendages.

The right forewings of 237 males and 98 females of *I. capreolus*, *I. cyane,* and morphologically intermediate individuals were used in geometric morphometric analyses (Table S1). Wings were removed and mounted on 25 × 75 mm microscope slides using Entellan® solution, covered with a 20 × 20 mm coverslip, and scanned at 1200 dpi using an Epson Perfection 2400 scanner. Caudal appendages of 201 males were photographed in lateral view using NIS-Elements BR (Instruments Nikon, 2014). Thirteen homologous landmarks were selected for forewings based on previously established landmark configurations (e.g., *Enallagma cyathigerum*: Bots et al., 2009), and 13 for caudal appendages (Fig. S1).

Images were digitized and landmarks were recorded in tpsDig2 v2.19 (Rohlf, 2015). Landmark configurations were subjected to Generalized Procrustes Analysis (GPA) separately for each structure to remove the effects of translation, rotation, and scale (Zelditch *et al*., 2012), producing Procrustes coordinates (shape variables) and Centroid Size (CS), an isometric estimator of structure size. Consensus configurations were generated for each structure and geographic group and visualized using deformation grids. Differences in CS among taxa (*I. capreolus*, *I. cyane*, and morphologically intermediate individuals) and geographic contexts (allopatry vs. sympatry) were evaluated using ANOVA followed by Tukey post hoc tests. Assumptions of normality and homoscedasticity were assessed using Shapiro–Wilk and Levene tests with Bonferroni correction (α = 0.05).

To evaluate morphological differentiation among taxa and geographic contexts (allopatry vs. sympatry), shape variation was analyzed using Canonical Variate Analysis (CVA) based on Procrustes coordinates, which maximizes differences among predefined groups relative to within-group variation. Significance of pairwise differences was assessed using permutation tests of Procrustes distances (10,000 permutations). Discriminant analyses were additionally conducted to evaluate shape differentiation among taxa and the assignment of individuals to parental or intermediate morphological categories. These analyses were performed in MorphoJ (Klingenberg, 2011). To evaluate classification accuracy, cross-validation analyses based on Mahalanobis distances were conducted in R 4.0.3 (R Core Team, 2013). Assignment of individuals to genetic clusters based on admixture coefficients (Q-values) is inherently sensitive to threshold choice, which may vary depending on the marker set and demographic context (Caniglia *et al*., 2020). Therefore, Q-values were interpreted conservatively as indicators of relative ancestry and admixture rather than as definitive assignments to hybrid classes (Pritchard *et al*., 2000; Sánchez-Guillén *et al*., 2011; Wellenreuther *et al*., 2018).

## RESULTS

### Species composition and temporal stability in sympatric localities

Both parental species (*I. capreolus* and *I. cyane*) and morphologically intermediate individuals were present in the sympatric localities of Anolaima and San Francisco, whereas El Colegio was characterized by the absence of *I. cyane* and the presence of *I. capreolus* and morphologically intermediate individuals (Fig. 1B). In El Colegio, *I. capreolus* was consistently the dominant taxon across sampling years, although morphologically intermediate individuals were also present at variable frequencies. In contrast, San Francisco was dominated by *I. cyane*, with a progressive increase in its relative frequency over time and a concomitant decline in the frequency of the remaining taxa. Anolaima showed a contrasting pattern, being consistently dominated by morphologically intermediate individuals, which represented the majority of individuals across sampling periods. Taxon composition differed significantly among localities (χ² = 849.87, df = 4, p < 0.001), confirming that each locality was characterized by a distinct assemblage dominated by different taxa.

### Genetic analyses of population structure and hybridization

#### Genetic diversity and population structure

DNA amplification and sequencing were successful for seven sampling localities, including allopatric and sympatric localities of *I. capreolus*, *I. cyane*, and morphologically intermediate individuals (Table S1). The mitochondrial markers analyzed were COI, Cytb, 16S, and ND1, and the nuclear markers were 28S, ITS, and PRMT. Sample sizes differed among loci due to variation in amplification success.

#### Genetic diversity, population genetic structure per marker

Genetic diversity varied among loci (Table S3). The mitochondrial markers COI and ND1 showed moderate haplotype diversity (Hd=0.470–0.640) and low nucleotide diversity (π=0.002–0.003). Cytb and 16S showed lower haplotype diversity (Hd=0.275–0.411) and nucleotide diversity values ranging from 0.001–0.002. For nuclear markers, ITS showed relatively high haplotype diversity (Hd=0.576) and nucleotide diversity (π=0.004), whereas PRMT showed the highest haplotype diversity (Hd=0.820) and nucleotide diversity (π=0.004). The 28S marker showed moderate haplotype diversity (Hd=0.516) and low nucleotide diversity (π=0.001).

AMOVA analyses revealed variation in the distribution of genetic variance among loci (Table S3). For mitochondrial markers, COI and ND1 showed a greater proportion of variation within populations (59–63%) than among populations (37–40%), with F_ST_ values ranging from 0.371 to 0.404 (p < 0.001). In contrast, Cytb and 16S showed a higher proportion of variation among populations (45–65%), with F_ST_ values ranging from 0.550 to 0.653 (p ≤ 0.020). For nuclear markers, ITS showed a more balanced distribution of genetic variance with 46% among populations and 54% within populations (F_ST_ = 0.460, p < 0.001). PRMT showed most of the variation within populations (86%), and a comparatively low F_ST_ value (0.140, p < 0.001). In contrast, 28S showed a high proportion of variation among populations (65%) together with a high F_ST_ value (0.651, p < 0.001).

#### Pairwise differentiation (F_ST_)

Pairwise F_ST_ values indicate that genetic differentiation between *I. capreolus* and *I. cyane* varied with geographic context.

In interspecific comparisons, differentiation was consistently higher in allopatric than in sympatric localities. Mitochondrial markers showed strong differentiation in allopatry (COI: F_ST_ = 0.546–0.727, p < 0.001; 16S: F_ST_ = 0.932, p < 0.001), whereas differentiation was lower in sympatry (COI: F_ST_ = 0.187, p < 0.001; 16S: F_ST_ = 0.230, p < 0.001).

Intraspecific comparisons revealed contrasting patterns between species. Within *I. capreolus*, differentiation remained high among allopatric populations (COI: F_ST_ = 0.576, p < 0.001), whereas comparisons involving sympatric localities showed lower or variable differentiation (COI: F_ST_ = 0.508–0.517, p < 0.00). 1In *I. cyane*, no significant differentiation was detected among localities in either allopatric or sympatric contexts. Comparisons involving morphologically intermediate individuals showed contrasting levels of differentiation relative to the parental species. For mitochondrial markers, morphologically intermediate individuals showed significant differentiation from *I. capreolus* localities in both allopatry and sympatry (COI: F_ST_ = 0.095–0.527, p < 0.001), whereas differentiation relative to *I. cyane* localities was lower and significant only in sympatry (COI: F_ST_ = 0.047, p < 0.001). Among nuclear markers, significant differentiation was detected only for 28S, which showed patterns similar to those observed for COI in interspecific comparisons and between *I. capreolus* and morphologically intermediate individuals from Anolaima (Table 2).

**Table 1.** Sampling sites of *Ischnura capreolus*, *I. cyane*, and putative hybrids from allopatric and sympatric localities.

| Species | Locality | Distribution | Latitude | Longitude | Altitude<br>(m a.s.l.) |
| --- | --- | --- | --- | --- | --- |
| *I. capreolus* | Saldaña, Tolima | Allopatric | 3°57'51.674''N | 72°52'20.332''W | 402 |
| *I. capreolus* | Puerto Nariño, Amazonas | Allopatric | 4°35'27,24''N | 70°13'4,8''W | 159 |
| *I. capreolus* | El Colegio, Cundinamarca | Sympatric | 4°32 .615'N | 74°26.243''W | 1507 |
| **Morphologically intermediate** | Anolaima, Cundinamarca | Sympatric | 5°37'1.6''N | 73°32'15.3''W | 1650 |
| *I. cyane* | San Francisco, Cundinamarca | Sympatric | 4°59.228'N | 74°17.107'W | 1627 |
| *I. cyane* | Villa de Leyva, Boyacá | Allopatric | 5°37'1.6''N | 73°32'15.3''W | 2172 |
| *I. cyane* | Barbosa, Santander | Allopatric | 4°45'43,74''N | 73°36'14,4''W | 1588 |

**Table 2.** Pairwise F_ST_ estimates for mitochondrial (16S, Cytb, COI, ND1) and nuclear (PRMT, 28S, ITS) markers among allopatric and sympatric localities of *Ischnura capreolus*, *I. cyane*, and putative hybrids. * Significant values (p < 0.05).

| Gene | Population | *I. cyane*<br>Cundinamarca | *I. cyane*<br>Boyacá | *I. cyane*<br>Santander | *I. capreolus*<br>Cundinamarca | *I. capreolus*<br>Tolima | *I. capreolus*<br>Amazonas | Morphologically<br>intermediate<br>Anolaima |
| --- | --- | --- | --- | --- | --- | --- | --- | --- |
| - | *I. cyane*<br>Cundinamarca | - |  |  |  |  |  |  |
| 16S | *I. cyane*<br>Boyacá | -0.06240 |  |  |  |  |  |  |
| Cytb |  | -0.01124 |  |  |  |  |  |  |
| COI |  | 0.00444 |  |  |  |  |  |  |
| ND1 |  | 0.18750 |  |  |  |  |  |  |
| PRMT |  | 0.08727 |  |  |  |  |  |  |
| 28S |  | -0.00645 |  |  |  |  |  |  |
| ITS |  | 0.00510 |  |  |  |  |  |  |
| 16S | *I. cyane*<br>Santander | - | - |  |  |  |  |  |
| Cytb |  | -0.08434 | 0.00000 |  |  |  |  |  |
| COI |  | 0.05660 | 0.04950 |  |  |  |  |  |
| ND1 |  | -0.07722 | 0.08623 |  |  |  |  |  |
| PRMT |  | - | - |  |  |  |  |  |
| 28S |  | -0.02612 | 0.01268 |  |  |  |  |  |
| ITS |  | -0.03355 | 0.01932 |  |  |  |  |  |
| 16S | *I. capreolus*<br>Cundinamarca | **0.23024\*** | 0.09856 | - |  |  |  |  |
| Cytb |  | 0.00000 | -0.01124 | -0.08434 |  |  |  |  |
| COI |  | **0.18681\*** | 0.14446 | 0.07454 |  |  |  |  |
| ND1 |  | -0.09615 | 0.30000 | -0.01867 |  |  |  |  |
| PRMT |  | 0.23395 | 0.21879 | - |  |  |  |  |
| 28S |  | **0.84137\*** | **0.92375\*** | **0.57745\*** |  |  |  |  |
| ITS |  | 0.57047 | 0.67923 | 0.56677 |  |  |  |  |
| 16S | *I. capreolus*<br>Tolima | **0.92743\*** | **0.93165\*** | - | **0.64383\*** |  |  |  |
| Cytb |  | 0.00454 | 0.00000 | -0.07784 | -0.04594 |  |  |  |
| COI |  | **0.76173\*** | **0.72700\*** | **0.56252\*** | **0.50839\*** |  |  |  |
| ND1 |  | 0.46885 | 0.47552 | 0.45569 | 0.48438 |  |  |  |
| PRMT |  | 0.25925 | 0.23878 | - | -0.02174 |  |  |  |
| 28S |  | **0.91977\*** | **1.00000\*** | **0.63462\*** | -0.00871 |  |  |  |
| ITS |  | 0.53983 | 0.63760 | 0.53079 | -0.11294 |  |  |  |
| 16S | ***I. capreolus***<br>**Amazonas** | - | - | - | - | - |  |  |
| Cytb |  | 0.97778 | 1 | 1 | 0.93407 | 0.95062 |  |  |
| COI |  | **0.74954\*** | **0.70128\*** | **0.54632\*** | **0.51754\*** | **0.57604\*** |  |  |
| ND1 |  | 0.64120 | 0.67673 | 0.64451 | 0.64394 | 0.33244 |  |  |
| PRMT |  | - | - | - | - | - |  |  |
| 28S |  | **0.90569\*** | **1.00000\*** | **0.59090\*** | -0.03226 | 0.00000 |  |  |
| ITS |  | 0.71288 | 0.86828 | 0.73880 | 0.33213 | 0.31124 |  |  |
| 16S | **Morphologically**<br>**intermediate**<br>**Anolaima** | 0.07194 | 0.00384 | - | 0.03033 | **0.81428\*** | - | - |
| Cytb |  | -0.07143 | -0.01124 | -0.08434 | 0.00000 | 0.00062 | 0.95604 |  |
| COI |  | **0.04667\*** | 0.05578 | 0.03837 | **0.09522\*** | **0.46188\*** | **0.52707\*** |  |
| ND1 |  | 0.03140 | -0.01083 | -0.02297 | 0.11433 | 0.36852 | 0.55416 |  |
| PRMT |  | 0.22260 | 0.21412 | - | 0.00698 | -0.01257 | - |  |
| 28S |  | 0.02765 | 0.18182 | -0.02833 | **0.64971\*** | **0.74507\*** | **0.70660\*** |  |
| ITS |  | 0.01252 | 0.08510 | 0.00395 | 0.48529 | 0.46739 | 0.60131 |  |

#### Admixture patterns and haplotype structure

STRUCTURE analyses of concatenated mitochondrial and nuclear datasets supported K=2 (ΔK=32.94) as the most likely number of genetic clusters according to the ΔK method of Evanno. Individuals of *I. capreolus* and *I. cyane* were primarily assigned to two distinct genetic clusters (green and red, respectively) (Fig. 2), but some individuals from sympatric localities (Anolaima, San Francisco, and El Colegio) showed mixed ancestry.

**Figure 2.**
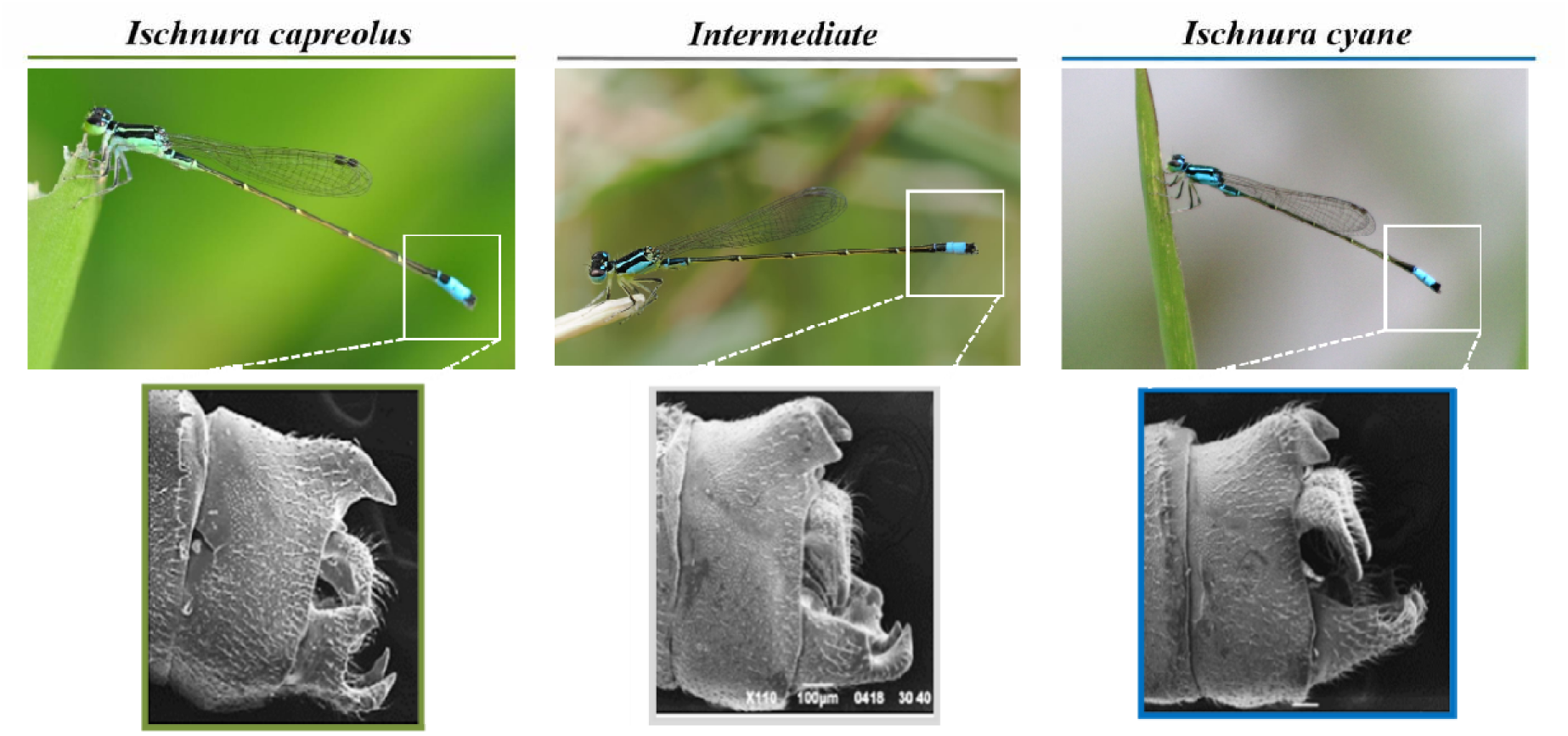
Morphometric variation of male caudal appendages. Representative variation in male caudal appendage morphology among *I. capreolus*, *I. cyane*, and morphologically intermediate individuals.

In several cases, individuals morphologically assigned to *I. capreolus* exhibited mitochondrial ancestry associated with *I. cyane* (Fig. S2). Nuclear loci generally showed stronger differentiation between clusters than mitochondrial loci (Fig. S3). When loci were analyzed separately, most markers also supported K=2, except ND1, which supported K=3. Most loci displayed bimodal genotype distributions, whereas PRMT showed *I. cyane* genotypes predominantly and very few admixed individuals.

Haplotype networks revealed species-specific haplotypes in allopatric populations of *I. capreolus* and *I. cyane*. In sympatric localities, several haplotypes were shared between species (Fig. 2). Mitochondrial loci (COI, Cytb, ND1 and 16S) were characterized by large central haplotypes containing individuals from multiple populations and both species, with several low-frequency haplotypes derived from them. In contrast, nuclear loci (particularly 28S and PRMT) showed haplotypes strongly structured by species. In the sympatric locality of Anolaima, individuals carried haplotypes associated with both parental species across several loci, whereas species-specific haplotypes were rare or absent. Individuals from this locality were primarily associated with haplotypes shared with either parental species rather than forming exclusive haplotype groups (Fig. S4, S5).

### Morphological differentiation and hybrid intermediacy: wing morphology

Thin-plate spline deformation grids further illustrated the wing shape differences between parental species and the intermediate individuals (Fig. S6). Procrustes ANOVA revealed significant differences in both wing size (CS) and wing shape (partial warps) among all taxon pairs *I. capreolus*, *I. cyane*, and morphologically intermediate individuals (CS: F_21,346_=369.69, p<0.0001; shape: F_44,7612_=97.06, p<0.0001). Significant differences were also detected between sexes (CS: F_1,347_=23.91, p<0.0001; shape: F_22,7634_=2.48, p=0.0001) and among localities (CS: F_6,342_=159.23, p<0.0001; shape: F_132,7524_=39.31, p<0.0001) (Table 3). Pairwise comparisons showed significant differences in both CS and wing shape among all taxon pairs (*I. cyane* vs. *I. capreolus*, *I. capreolus* and morphologically intermediate individuals) (Table S4, S6). Among localities, 14 of 21 comparisons were significant for CS whereas 20 of 21 comparisons were significant for wing shape (Table S5, S7; Fig. 4).

**Figure 3.**
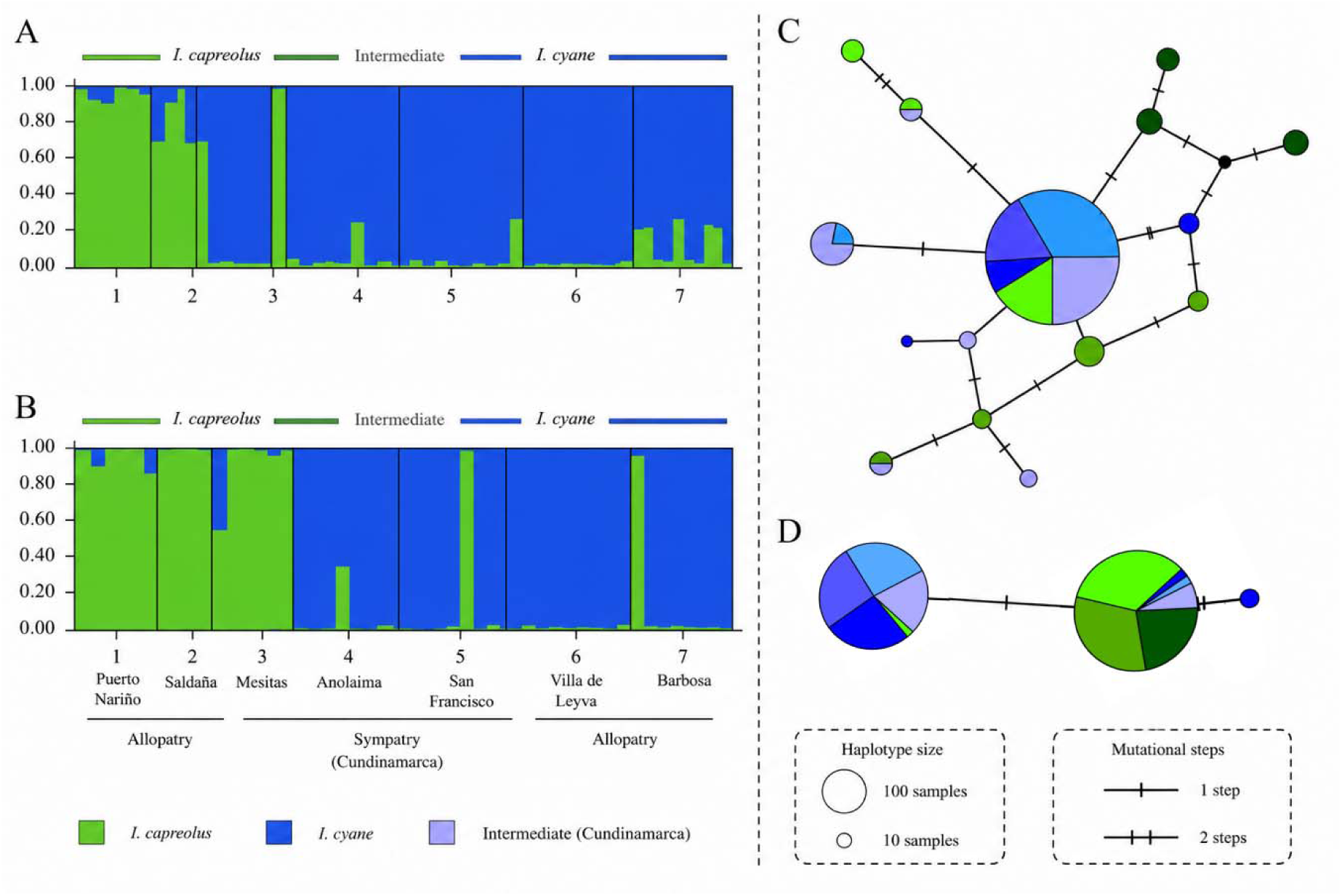
Genetic structure and haplotype networks. Bayesian clustering analyses performed with STRUCTURE based on concatenated loci for *I. capreolus*, *I. cyane*, and morphologically intermediate individuals from allopatric and sympatric localities. Each vertical bar represents an individual, and colors indicate the proportional membership (Q) to each genetic cluster (K=2). Individuals are grouped by locality: 1. Puerto Nariño (allopatry, *I. capreolus*), 2. Saldaña (allopatry, *I. capreolus*), 3. El Colegio (sympatry), 4. Anolaima (sympatry), 5. San Francisco (sympatry), 6. Villa de Leyva (allopatry, *I. cyane*), and 7. Barbosa (*I. cyane*). **(A)** Mitochondrial loci (COI–ND1). **(B)** Nuclear loci (ITS–28S). **(C)** Nuclear 28S haplotype network showing two predominant haplotypes largely associated with *I. capreolus* and *I. cyane*, respectively, with morphologically intermediate individuals occurring within both groups. Compared with mitochondrial loci, the 28S marker exhibits clearer species differentiation and limited haplotype sharing in sympatric localities. **(D)** Mitochondrial COI haplotype network dominated by a large central haplotype shared among individuals from multiple localities and both species, with several low-frequency derived haplotypes radiating from it. Individuals from sympatric localities, including morphologically intermediate individuals from Anolaima, share haplotypes, indicating haplotype mixing and introgression in the contact zone.

**Figure 4.**
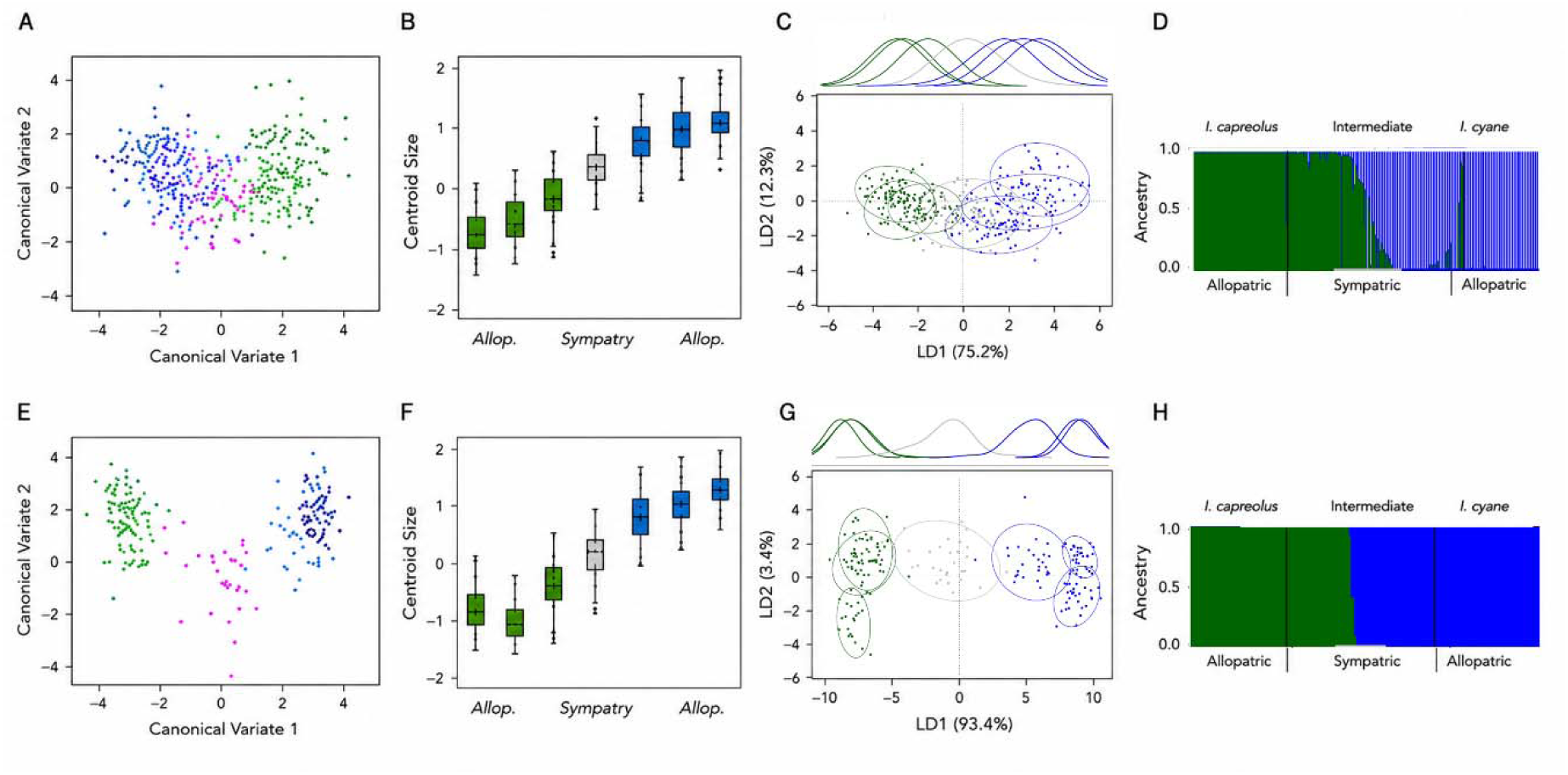
Morphometric differentiation and intermediacy. **(A)** Canonical variate analysis (CVA) of wing shape and **(B)** centroid size variation of wings for individuals from allopatric populations of *I. capreolus* (dark and light green), sympatric populations including morphologically intermediate individuals (grey and magenta), and allopatric populations of *I. cyane* (light and dark blue). **(C)** Linear discriminant analysis (LDA) of wing morphology showing partial overlap among sympatric intermediate individuals and parental species. LD1 and LD2 explain 75.2% and 12.3% of the variation, respectively. **(D)** Bayesian clustering analysis based on wing morphology showing admixture patterns between *I. capreolus*, intermediate individuals, and *I. cyane* across allopatric and sympatric populations. **(E)** Canonical variate analysis (CVA) of caudal appendage shape and **(F)** centroid size variation of caudal appendages. **(G)** Linear discriminant analysis (LDA) of caudal appendage morphology showing clearer differentiation between parental species and intermediate placement of sympatric individuals. LD1 and LD2 explain 93.4% and 3.4% of the variation, respectively. **(H)** Bayesian clustering analysis based on caudal appendage morphology, showing strong assignment of individuals to parental morphotypes and reduced admixture compared with wing morphology. Colors indicate *I. capreolus* (green), morphologically intermediate individuals from sympatry (grey/magenta), and *I. cyane* (blue).

**Table 3.** Procrustes ANOVA of centroid size and shape of the forewings of *I. capreolus, I. cyane* and putative hybrids: (SS) Sum of squares; (MS) Mean squares, measured in Procrustes distances. Taxa (*I. capreolus*, *I. cyane*, and putative hybrids). Sex (males and females). Localities (Saldaña, Puerto Nariño, El Colegio, Anolaima, San Francisco, Villa de Leyva, and Barbosa).

| Centroid size | SS | MS | Df | F | Valor P |
| --- | --- | --- | --- | --- | --- |
| Taxa | 2155912 | 1077956 | 21 | 369.69 | <0.0001 |
| Residual | 1009134 | 2917 | 346 |  |  |
| Sex | 203999 | 203999 | 1 | 23.91 | <0.0001 |
| Residual | 2961047 | 8533 | 347 |  |  |
| Locality | 2330721 | 388453 | 6 | 159.23 | <0.0001 |
| Residual | 834324 | 2440 | 342 |  |  |
| Shape | SS | MS | Df | F | Valor P |
| Taxa | 0.219 | 0.005 | 44 | 97.06 | <0.0001 |
| Residual | 0.391 | □ 0.001 | 7612 |  |  |
| Sex | 0.004 | □ 0.001 | 22 | 2.48 | 0.0001 |
| Residual | 0.606 | □ 0.001 | 7634 |  |  |
| Locality | 0.249 | 0.002 | 132 | 39.31 | <0.0001 |
| Residual | 0.361 | □ 0.001 | 7524 |  |  |

Canonical Variate Analysis revealed differentiation in wing shape among the three taxa in both allopatric and sympatric contexts (Fig. 4). Clustering patterns showed clearer separation among individuals from allopatric populations, whereas individuals from sympatric localities showed greater overlap. Discriminant analyses based on wing shape separated *I. capreolus* and *I. cyane* in allopatric populations, whereas individuals from sympatric localities overlapped with both taxa in morphospace.

Additionally, cross-validation analyses showed clear assignment of individuals to parental species in allopatric populations, with all individuals correctly classified except one male from Saldaña. In sympatric localities, individuals were assigned to both parental and morphologically intermediate individuals. In Anolaima, individuals were distributed among pure *I. capreolus*, pure *I. cyane*, and morphologically intermediate individuals. In El Colegio, most individuals were assigned to pure *I. capreolus*, with a smaller number assigned to morphologically intermediate individuals. In San Francisco, most individuals were assigned to pure *I. cyane*, with a small number assigned to morphologically intermediate individuals (Fig. 4).

### Morphological differentiation and hybrid intermediacy: caudal appendages

Thin-plate spline deformation grids further illustrated the shape differences in caudal appendages between parental species and the intermediate individuals (Fig. S7). Procrustes ANOVA showed significant differences in both CS and shape (partial warps) of male caudal appendages among *I. capreolus*, *I. cyane*, and morphologically intermediate individuals (size: F_2,197_=379.87, p<0.0001; shape: F_44,4334_=166.67, p<0.0001). Significant differences were also detected among localities (size: F_6,193_=185.66, p<0.0001; shape: F_132,4246_=68.37, p<0.0001) (Table 4).

**Table 4.** Procrustes ANOVA of centroid size and shape of the male caudal appendages of *I. capreolus*, *I. cyane,* and putative hybrids: (SS) Sum of squares; (MS) Mean squares, measured in Procrustes distances. Taxa (*I. capreolus*, *I. cyane*, and putative hybrids). Localities (Puerto Nariño, Saldaña, El Colegio, Anolaima, San Francisco, Villa de Leyva, and Barbosa).

| Centroid size | SS | MS | Df | F | Valor P |
| --- | --- | --- | --- | --- | --- |
| Taxa | 1880056 | 940028 | 2 | 379.87 | <0.0001 |
| Residual | 487495 | 2475 | 197 |  |  |
| Locality | 2017937 | 336323 | 6 | 185.66 | <0.0001 |
| Residual | 349614 | 1811 | 193 |  |  |
| Shape | SS | MS | Df | F | Valor P |
| Taxa | 2.286 | 0.052 | 44 | 166.67 | <0.0001 |
| Residual | 1.351 | □ 0.001 | 4334 |  |  |
| Locality | 2.474 | 0.019 | 132 | 68.37 | <0.0001 |
| Residual | 1.164 | □ 0.001 | 4246 |  |  |

Pairwise comparisons were significant in both CS and shape among all taxon pairs (Table S8, S10). Among localities, 6 of 21 comparisons were significant for CS, whereas 19 of 21 comparisons were significant for caudal appendage shape (Table S9, S11). Canonical Variate Analysis revealed differentiation in caudal appendage shape among the three taxa (Fig. 4). Separation among taxa was evident in both allopatric and sympatric contexts, although clustering patterns showed greater separation in allopatric populations. Discriminant analyses revealed clear separation between *I. capreolus* and *I. cyane* in allopatric populations, whereas individuals from Anolaima overlapped with both parental species in sympatry (Fig. 4).

Cross-validation analyses showed complete assignment of males to parental species in allopatric populations. In sympatric populations, most individuals were assigned to parental categories, although some individuals occupied intermediate positions in morphospace. A small number of individuals from Anolaima occupied intermediate positions between parental categories (Fig. 4).

## DISCUSSION

Hybridization is increasingly recognized as a widespread and evolutionarily significant process across the tree of life, rather than a rare or purely maladaptive phenomenon (Peñalba *et al*., 2024). By promoting gene flow between species, hybridization can reshape species boundaries and redistribute genetic variation across populations, often generating admixed individuals with intermediate or transgressive phenotypes. Our results provide evidence of hybridization between *I. capreolus* and *I. cyane* and show that sympatric localities, particularly Anolaima, are characterized by a high frequency of morphologically intermediate individuals with admixed genetic signatures, indicating that these individuals represent admixed forms derived from hybridization between the two species. The persistence of these admixed individuals across sympatric localities and sampling periods suggests that hybridization in this system is not limited to rare or transient events (Fig. 5).

**Figure 5.**
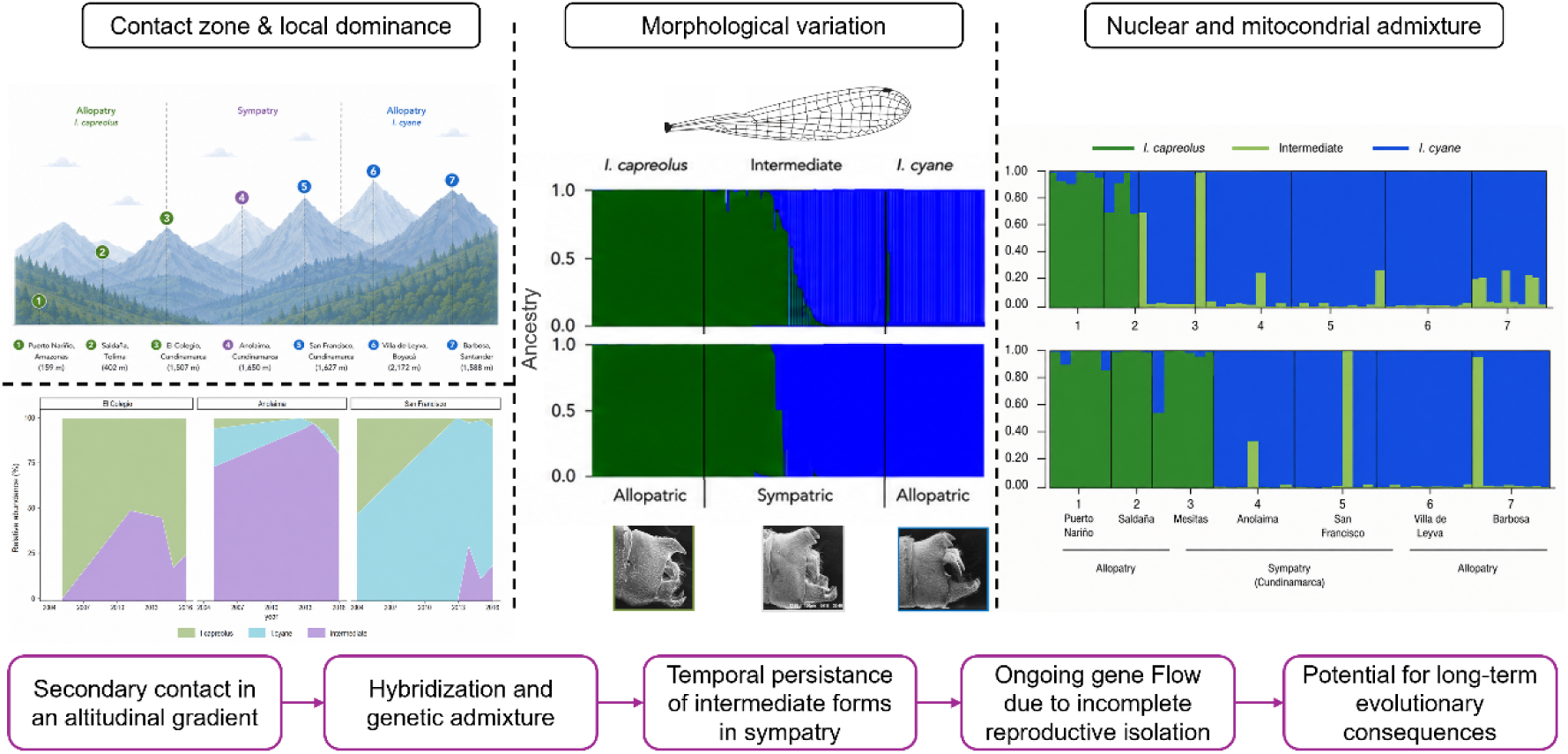
Proposed scenario of secondary contact and hybridization between *I. capreolus* and *I. cyane*. Geographic overlap in an altitudinal contact zone promotes hybridization and genetic admixture, generating morphologically intermediate individuals that persist through time in sympatric populations. Patterns of mitochondrial and nuclear admixture indicate ongoing gene flow and incomplete reproductive isolation, suggesting potential long-term evolutionary consequences for both species.

This combination of admixture and phenotypic intermediacy is consistent with our initial predictions and aligns with previous studies showing that hybridization can generate predictable patterns of genetic and phenotypic intermediacy (Abbott et al., 2013; Monetti et al., 2002; Rieseberg et al., 2007). In this system, both genetic and morphological datasets independently identified intermediate individuals and reduced differentiation in sympatric localities. These concordant patterns support the interpretation that these forms represent hybrids rather than phenotypic variants. This concordance between datasets is particularly important given that hybrid identification based solely on morphology can be misleading due to the occurrence of transgressive or parent-like traits in later-generation hybrids (Dittrich-Reed & Fitzpatrick, 2013).

Genetic analyses provide consistent evidence of admixture between *I. capreolus* and *I. cyane*, across sympatric localities, where reduced differentiation and haplotype sharing indicate ongoing or recent gene flow. The contrast between clear genetic differentiation in allopatric populations and admixture in sympatry suggests that reproductive barriers are incomplete. In addition, the occurrence of mito-nuclear discordance is consistent with asymmetric introgression, a pattern commonly associated with unidirectional hybridization or differential permeability of the genome to gene flow. Asymmetric reproductive isolation has been frequently reported in odonates, where mating interactions and hybridization rates differ between reciprocal crosses (Tynkkynen *et al*., 2008b; Sánchez-Guillén *et al*., 2012, 2014b; Barnard *et al*., 2017; Solano *et al*., 2018), suggesting that gene flow between species may often be directionally biased.

In addition, the differences observed among genetic markers provide further insight into the dynamics of hybridization in this system. Patterns of genetic differentiation (F_ST_), haplotype structure, and admixture varied among loci, revealing a heterogeneous genomic response to gene flow. While some markers (e.g., 16S and 28S) retained higher levels of population differentiation and more structured haplotype distributions, others (e.g., COI and PRMT) showed lower differentiation, greater haplotype sharing, and higher levels of admixture. This contrast suggests that different loci vary in their permeability to introgression. Markers showing lower differentiation and greater haplotype sharing are likely capturing ongoing gene flow, whereas those retaining stronger structure may reflect genomic regions more resistant to introgression, potentially due to selective constraints or differences in inheritance, or stochastic lineage sorting. This locus-specific signal helps explain the combined patterns of admixture and differentiation observed in sympatric localities and reinforces the interpretation of incomplete reproductive isolation between species, a pattern commonly observed in hybrid zones and consistent with differential introgression across the genome (Harrison & Larson, 2014; Swaegers *et al*., 2022).

Patterns of morphological variation further support this interpretation but also reveal important differences between wing morphology and reproductive structures. Wing morphology showed extensive overlap and intermediacy in sympatric localities, consistent with reduced differentiation under gene flow. This is particularly relevant given that wing morphometry has been widely proposed as a reliable tool for species identification (Sontigun *et al*., 2017), as wing structure plays a key role in functions such as predator avoidance, foraging, and courtship behavior, all of which depend on efficient and precise flight performance (Bots *et al*., 2009). Wing size and shape differed between *I. capreolus* and *I. cyane*, with females showing greater differentiation than males, particularly in allopatric populations. This differentiation was reduced in sympatry, consistent with introgression between species. In contrast, reproductive traits such as male caudal appendages, although also showing some degree of intermediacy, retained clearer differentiation in allopatric populations and a more structured distribution overall. This contrast suggests that different components of the phenotype respond unequally to hybridization, potentially reflecting differences in the selective pressures acting on them. In odonates, male caudal appendages play a central role in species recognition and mating compatibility and are therefore expected to be more strongly associated with reproductive isolation (Paulson, 1974; Wellenreuther & Sánchez Guillén, 2015; Barnard *et al*., 2017; Ordaz-Morales *et al*., 2026). However, the occurrence of intermediate genital morphology in sympatric localities suggests that these traits do not constitute a complete reproductive barrier in this system, allowing hybridization despite clear morphological differentiation between species.

The combination of genetic admixture and phenotypic intermediacy, together with the high frequency of intermediate individuals, is consistent with a hybrid zone characterized by ongoing gene flow. This contrasts with classical tension zones, typically maintained by selection against hybrids and characterized by narrow spatial structure and low hybrid frequencies (Barton & Hewitt, 1985; Harrison, 1990), suggesting that hybridization in this system persists despite the maintenance of species differences. Such conditions, in which hybrids are frequent and persist over time, are expected to facilitate the maintenance and potential stabilization of admixed populations. In this context, hybridization may extend beyond transient introgression and contribute to the long-term maintenance of admixed populations. Although hybrid speciation remains difficult to demonstrate conclusively, especially in animals, it is increasingly recognized as a possible outcome under specific evolutionary conditions (Abbott *et al*., 2013; Peñalba *et al*., 2024). Accordingly, the patterns observed here are consistent with expectations for systems in which hybridization may lead to the establishment of stable admixed populations, and potentially to early stages of lineage differentiation. Whether this pattern reflects a transient hybrid swarm or the early stages of a hybrid lineage remains unresolved. However, the consistency of genetic and morphological signals, together with the high frequency of intermediate individuals in sympatry, highlights the potential for hybridization to contribute to the generation and maintenance of evolutionary novelty in this system.

### Conclusions

Hybridization between *I. capreolus* and *I. cyane* is supported by consistent patterns of genetic admixture and morphological intermediacy in sympatric localities. These results highlight the role of hybridization as a dynamic process shaping genetic and phenotypic variation, particularly in systems where reproductive barriers are incomplete. The persistence and spatial structuring of intermediate forms suggest that hybridization may contribute not only to gene flow between species, but also to the generation of persistent admixed populations and potentially novel evolutionary trajectories.

## Notes

### Competing Interest Statement

The authors have declared no competing interest.

### Funding

- Universidad de Los Andes
- Secretaría de Ciencia, Humanidades, Tecnología e Innovación: 282922
- Ministry of Science, Innovation and Universities, Spain: PGC2018-096656-B-I00, PID2023-147268NB-I00

