## Supplementary File for "Temporal persistence of admixed phenotypes in a damselfly hybrid zone"

### Supplementary files

**Table S1.** Number of wings and male appendages analyzed, and the number of individuals sequenced for mitochondrial and nuclear markers of *Ischnura capreolus*, *I. cyane*, and putative hybrids.

| Species | Mitochondrial | | | | Nuclear | | | Wings | | Caudal appendages |
|---|---|---|---|---|---|---|---|---|---|---|
| | 16S | COI | Cytb | ND1 | 28S | ITS | PRMT | N ♂ | N ♀ | N ♂ |
| ***I. capreolus*** | 20 | 7 | 9 | 11 | 14 | 7 | 13 | 40 | 24 | 35 |
| ***I. capreolus*** | - | 8 | 1 | 6 | 10 | 7 | - | 19 | 9 | 19 |
| ***I. capreolus*** | 15 | 13 | 10 | 9 | 16 | 7 | 16 | 30 | 15 | 29 |
| **Putative hybrids** | 33 | 32 | 10 | 11 | 12 | 12 | 12 | 47 | 16 | 31 |
| ***I. cyane*** | 31 | 30 | 10 | 9 | 13 | 14 | 14 | 42 | 17 | 29 |
| ***I. cyane*** | 8 | 16 | 9 | 9 | 12 | 15 | 7 | 29 | 9 | 28 |
| ***I. cyane*** | - | 10 | 5 | 12 | 14 | 12 | - | 30 | 8 | 30 |

**Table S2.** Primers used for amplification of mitochondrial and nuclear markers used in *I. capreolus* and *I. cyane* from Colombia.

| Gen | Forward y Reverse | N | Pb | Annealing T (°C) | Reference |
|---|---|---|---|---|---|
| COI | F:5´-GGTCAACAAATCATAAAGATATTGG-3´<br>R:5´-TCAGGGTGACCAAAAAATCA -3´ | 116 | 650 | 45° | (Folmer, Hoeh et al. 1994) |
| Cytb | F:5´-TATGTACTACCATGAGGACAAATATC-3´<br>R:5´-TATTTCTTTATTATGTTTTCAAAAC-3´ | 54 | 520 | 48° | (Simon, Frati et al. 1994) |
| ND1 | F:5´-ACATGAATTGGAGCTCGACCAGT-3´<br>R:5´-GATTTTGCTGAAGGTGAATCAGA-3´ | 67 | 540 | 48° | (Simon, Frati et al. 1994) |
| 16S | F:5´-CCGGTTTGAACTCAGATCACGT-3´<br>R:5´-CGCCTGTTTAACAAAAACAT-3´ | 107 | 500 | 44° | (Simon, Frati et al. 1994) |
| ITS | F:5´-CTTTGTACACACCGCCCGTCGCT-3´<br>R:5´-TTTCACTCGCCGTTACTAAGGGAATC-3´ | 62 | 740 | 52° | Modificados por (Elbadri, De Ley et al. 2002) |
| 28S | F:5´-AAGGTAGCCAAATGCCTCATC-3´<br>R:5´-AGTAGGGTAAAACTAACCT-3´ | 91 | 794 | 49° | (Hillis and Dixon 1991) |
| PRMT | F:5̓ TCGACTCGTATGCGCATTTC-3´<br>R: 5̓ TGCCACCTTCCTAATAGAGC-3´ | 74 | 750 | 55° | (Ferreira, Lorenzo-Carballa et al. 2014) |

**Table S3.** Genetic diversity indices and AMOVA-based population differentiation for mitochondrial and nuclear markers. **N** = number of sequences; **Length** = analyzed fragment length; **S** = number of polymorphic sites; **h** = number of haplotypes; **Hd** = haplotype diversity; **π** = nucleotide diversity. **$F_{ST}$** and **p-values** were assessed using permutation tests. Genetic variation based on **$F_{ST}$** is expressed as the proportion partitioned **among** and **within populations**.

| Gene | N | Length (bp) | S | h | Hd | π | $F_{ST}$ | p-value | Among populations (%) | Within populations (%) |
|---|---|---|---|---|---|---|---|---|---|---|
| **COI – mtDNA** | 116 | 568 | 12 | 14 | 0.470 | 0.00165 | **0.371** | <0.001 | 37.12 | 62.88 |
| **ND1 – mtDNA** | 67 | 378 | 8 | 10 | 0.637 | 0.00321 | **0.404** | <0.001 | 40.37 | 59.63 |
| **Cytb – mtDNA** | 54 | 325 | 13 | 7 | 0.275 | 0.00192 | **0.55** | 0.02 | 55.04 | 44.96 |
| **16S – mtDNA** | 108 | 431 | 4 | 5 | 0.411 | 0.00103 | **0.653** | <0.001 | 65.32 | 34.68 |
| **28S – nuclear** | 92 | 746 | 5 | 3 | 0.516 | 0.00079 | **0.651** | <0.001 | 65.12 | 34.88 |
| **ITS – nuclear** | 74 | 412 | 9 | 19 | 0.576 | 0.00404 | **0.46** | <0.001 | 46.00 | 54.00 |
| **PRMT – nuclear** | 62 | 431 | 23 | 20 | 0.820 | 0.00401 | **0.14** | <0.001 | 14.00 | 86.00 |

**Table S4.** *Post hoc* Tukey tests of centroid size of the forewings. Pairwise comparisons between taxa: *I. capreolus*. *I. cyane* and putative hybrids.

| A | ***I. capreolus*** | ***I. cyane*** | **Hybrids** |
|---|---|---|---|
| ***I. capreolus*** | - | <0.001 | <0.001 |
| ***I. cyane*** | **7.9** | - | <0.001 |
| **Hybrids** | **14.3** | **22.3** | - |

**Table S5.** *Post hoc* Tukey tests of centroid size of the forewings. Pairwise comparisons between populations: Saldaña, Puerto Nariño, Mesitas, Anolaima, San Francisco, Villa de Leyva and Barbosa.

| B | **Puerto Nariño** | **Saldaña** | **Mesitas** | **Anolaima** | **San Francisco** | **Barbosa** | **Villa de Leyva** |
|---|---|---|---|---|---|---|---|
| **Puerto Nariño** | - | <0.001 | <0.001 | <0.001 | <0.001 | 0.003 | 0.001 |
| **Saldaña** | **7.4** | - | 0.959 | <0.001 | 0.001 | 0.771 | 0.905 |
| **Mesitas** | **6.1** | 1.4 | - | <0.001 | <0.001 | 0.999 | 1.000 |
| **Anolaima** | **13.8** | **6.3** | **7.7** | - | 0.999 | <0.001 | <0.001 |
| **San Francisco** | **13.1** | **5.6** | **7.0** | 0.7 | - | <0.001 | <0.001 |
| **Barbosa** | **5.4** | 2.1 | 0.7 | **8.4** | **7.7** | - | <0.001 |
| **Villa de Leyva** | **5.8** | 1.7 | 0.3 | **8.0** | **7.3** | 0.4 | - |

**Table S6.** *Post hoc* Turkey tests of shape of the forewings. Pairwise comparisons between taxa: *I. capreolus*. *I. cyane* and putative hybrids.

| A | *I. capreolus* | *I. cyane* | Hybrids |
|---|---|---|---|
| ***I. capreolus*** | - | <0.001 | <0.001 |
| ***I. cyane*** | **2.3** | - | <0.001 |
| **Hybrids** | **0.2** | **2.2** | - |

**Table S7.** *Post hoc* Turkey tests of shape of the forewings. Pairwise comparisons between populations: Saldaña, Puerto Nariño, Mesitas, Anolaima, San Francisco, Villa de Leyva and Barbosa.

| B | Puerto Nariño | Saldaña | Mesitas | Anolaima | San Francisco | Barbosa | Villa de Leyva |
|---|---|---|---|---|---|---|---|
| **Puerto Nariño** | - | 0.150 | <0.001 | <0.001 | <0.001 | <0.001 | <0.001 |
| **Saldaña** | 1.3 | - | <0.001 | <0.001 | <0.001 | <0.001 | <0.001 |
| **Mesitas** | **1.4** | **0.1** | - | <0.001 | <0.001 | <0.001 | <0.001 |
| **Anolaima** | **2.8** | **1.5** | **1.4** | - | <0.001 | <0.001 | <0.001 |
| **San Francisco** | **2.5** | **1.2** | **1.1** | **0.3** | - | 0.025 | <0.001 |
| **Barbosa** | **2.6** | **1.2** | **1.1** | **0.3** | **0.4** | - | <0.001 |
| **Villa de Leyva** | **3.2** | **1.8** | **1.7** | **0.3** | **0.6** | **0.6** | - |

**Table S8.** *Post hoc* Turkey tests of centroid size of the male caudal appendages. Pairwise comparisons between taxa: *I. capreolus*, *I. cyane*, and putative hybrids.

| Species | *I. capreolus* | *I. cyane* | Hybrids |
|---|---|---|---|
| ***I. capreolus*** | - | <0.001 | <0.001 |
| ***I. cyane*** | **7.866** | - | <0.001 |
| **Hybrids** | **16.04** | **23.91** | - |

**Table S9.** *Post hoc* Turkey tests of centroid size of the male caudal appendages. Pairwise comparisons between populations: Puerto Nariño, Saldaña, Mesitas, Anolaima, San Francisco, Villa de Leyva, and Barbosa.

| Locality | Puerto Nariño | Saldaña | Mesitas | Anolaima | San Francisco | Barbosa | Villa de Leyva |
|---|---|---|---|---|---|---|---|
| Puerto Nariño | | <0.001 | 0.041 | 0.003 | <0.001 | <0.001 | < 0.001 |
| Saldaña | **6.650** | | 0.626 | 0.971 | 1.000 | 0.999 | 0.998 |
| Mesitas | **4.270** | 2.383 | | 0.988 | 0.853 | 0.289 | 0.270 |
| Anolaima | **5.360** | 1.291 | 1.092 | | 0.999 | 0.780 | 0.760 |
| San Francisco | **6.110** | 0.546 | 1.837 | 0.745 | | 0.971 | 0.965 |
| Barbosa | **7.400** | 0.745 | 3.128 | 2.036 | 1.291 | | 1.000 |
| Villa de Leyva | **7.450** | 0.794 | 3.178 | 2.085 | 1.341 | 0.050 | |

**Table S10.** *Post hoc* Turkey tests of shape of the male caudal appendages. Pairwise comparisons between taxa: *I. capreolus*. *I. cyane* and putative hybrids.

| Species | *I. capreolus* | *I. cyane* | Hybrids |
|---|---|---|---|
| ***I. capreolus*** | - | <0.001 | <0.001 |
| ***I. cyane*** | **6.5** | - | <0.001 |
| **Hybrids** | **0.5** | **7** | - |

**Table S11.** *Post hoc* Turkey tests of shape of the male caudal appendages. Pairwise comparisons between populations: Puerto Nariño, Saldaña, Mesitas, Anolaima, San Francisco de Sales and Villa de Leyva and Barbosa.

| Locality | Puerto Nariño | Saldaña | Mesitas | Anolaima | San Francisco | Barbosa | Villa de Leyva |
|---|---|---|---|---|---|---|---|
| Puerto Nariño | - | 0.100 | <0.001 | <0.001 | <0.001 | <0.001 | <0.001 |
| Saldaña | 4.6 | - | 0.765 | <0.001 | <0.001 | <0.001 | <0.001 |
| Mesitas | **6.3** | 1.7 | - | <0.001 | <0.001 | <0.001 | <0.001 |
| Anolaima | **7.2** | **2.7** | **0.9** | - | <0.001 | <0.001 | <0.001 |
| San Francisco | **8.1** | **3.5** | **1.8** | **0.9** | - | <0.001 | <0.001 |
| Barbosa | **6.4** | **1.8** | **0.1** | **0.8** | **1.7** | - | 0.211 |
| Villa de Leyva | **8.3** | **3.8** | **2.0** | **1.1** | **0.2** | 1.9 | - |

**Figure S1. Morphometric landmarks.** **(A)** Right forewing landmarks (n = 13) selected for geometric morphometric analyses of *I. capreolus*, *I. cyane*, and morphologically intermediate individuals. These landmarks were used to estimate wing centroid size (CS) and shape variation (partial warps). Landmark positions on the right forewing of *Ischnura* correspond to: (1) second antenodal vein; (2) birth RP; (3) intersection AA2; (4) distal edge MP; (5) distal edge RP3–4; (6) distal border RP2; (7) distal border RP1; (8) distal anterior border of the pterostigma; (9) proximal anterior border of the pterostigma; (10) beginning IR1; (11) beginning RP2 birth; (12) beginning RP3–4; and (13) node intersection. Wing venation terminology follows Riek and Kukalová-Peck (1984). **(B)** Male caudal appendage landmarks (n = 13) selected for geometric morphometric analyses of *I. capreolus*, *I. cyane*, and morphologically intermediate individuals. Landmarks correspond to: (1) base of the tenth abdominal segment; (2) medial angle of the tenth abdominal segment; (3) caudal tip of the tenth abdominal segment; (4) medial angle of the tenth segment; (5) medial angle of the tenth segment connected to the cerci base; (6) base of the cerci; (7) medial angle of the cerci; (8) base of the upper ventral branch of the paraproct; (9) medial angle between paraproct branches; (10) tip of the lower ventral branch of the paraproct; (11) angle of the lower paraproct branch; (12) lower base of the paraproct; and (13) base of the tenth abdominal segment.

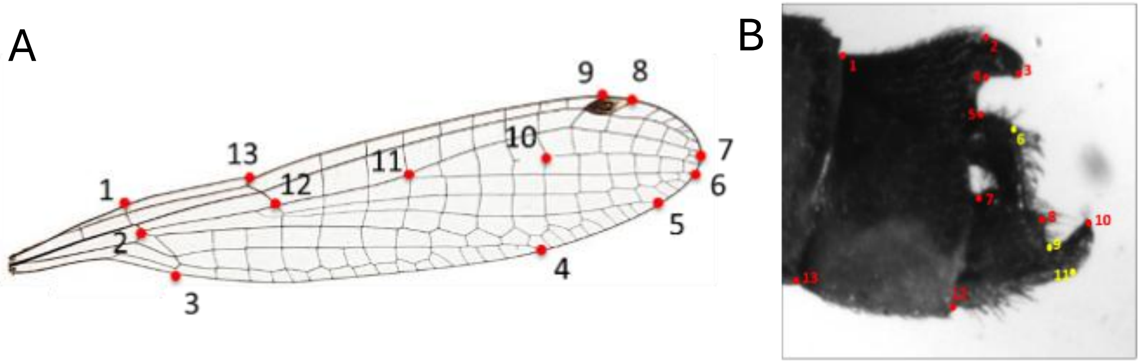

**Figure S2.** STRUCTURE assignment plots for mitochondrial genes in *I. capreolus*, *I. cyane* and putative hybrids from Colombia. Sampling localities are: 1 = Puerto Nariño (Amazonas), 2 = Saldaña (Tolima), 3 = Mesitas (Cundinamarca), 4 = Anolaima (Cundinamarca), 5 = San Francisco (Cundinamarca), 6 = Villa de Leyva (Boyacá), and 7 = Barbosa (Santander). Panels correspond to individual loci: **A)** 16S, **B)** COI, **C)** Cytb, and **D)** ND1.

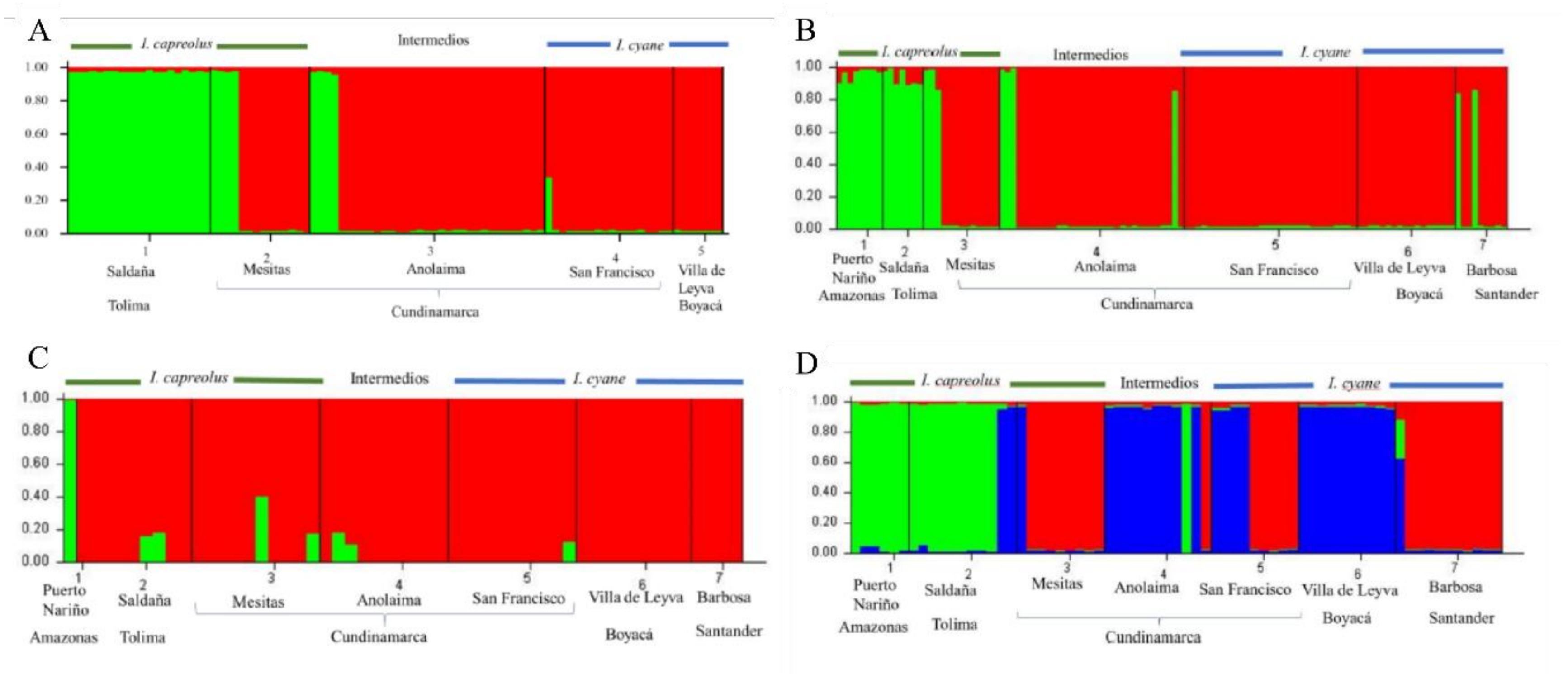

**Figure S3.** STRUCTURE assignment plots for nuclear genes in *I. capreolus*, *I. cyane* and putative hybrids from Colombia. Sampling localities are: 1 = Puerto Nariño (Amazonas), 2 = Saldaña (Tolima), 3 = Mesitas (Cundinamarca), 4 = Anolaima (Cundinamarca), 5 = San Francisco (Cundinamarca), 6 = Villa de Leyva (Boyacá), and 7 = Barbosa (Santander). Panels correspond to individual loci: **A) 28**S, **B)** ITS, and **C)** PMRT.

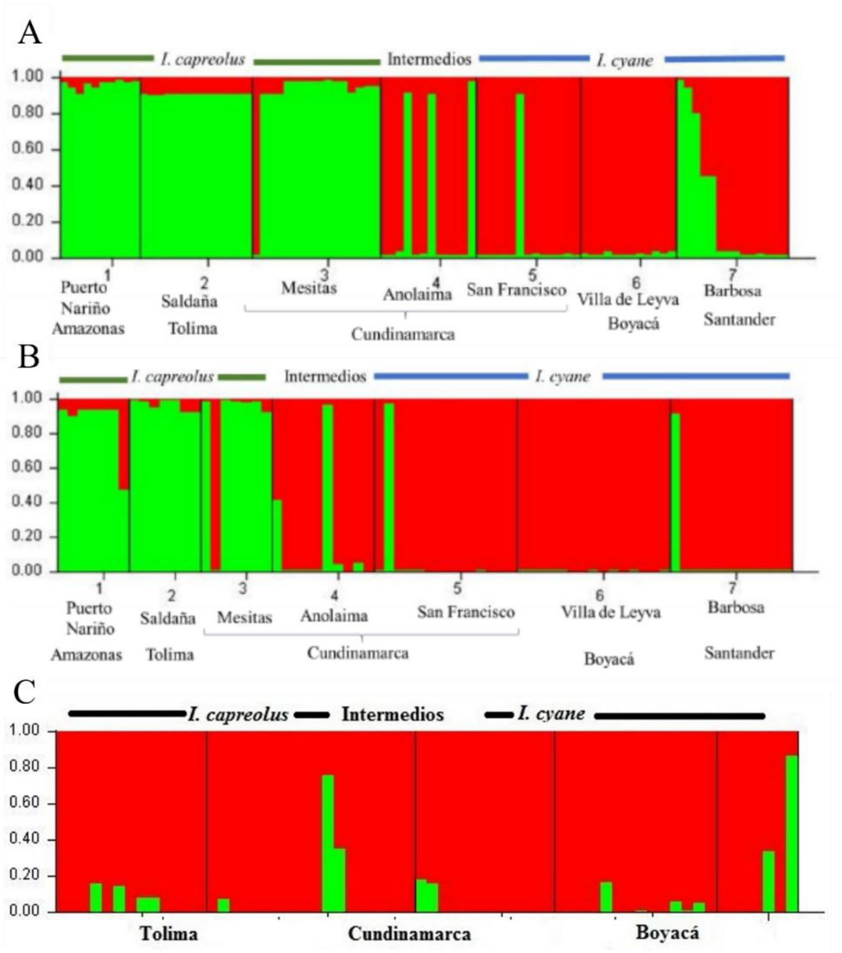

**Figure S4.** Haplotype networks reconstructed for mitochondrial genes in *I. capreolus*, *I. cyane*, and putative hybrids sampled across allopatric and sympatric populations. Networks are shown for **A)** 16S, **B)** Cytb, and **C)** ND1.

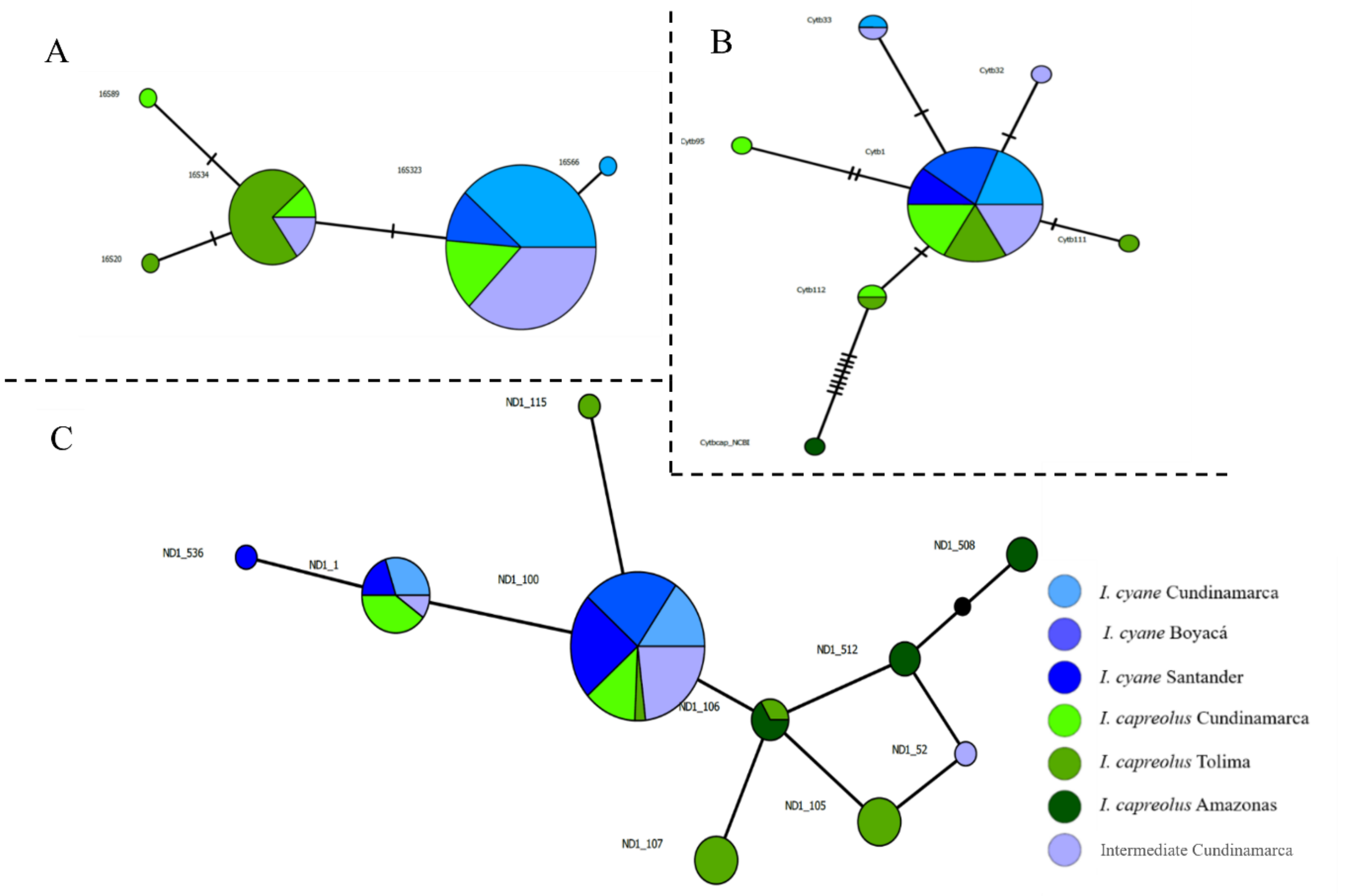

**Figure S5.** Haplotype networks reconstructed for nuclear genes in *I. capreolus*, *I. cyane*, and putative hybrids sampled across allopatric and sympatric populations. Networks are shown for **A)** ITS and **B)** PMRT.

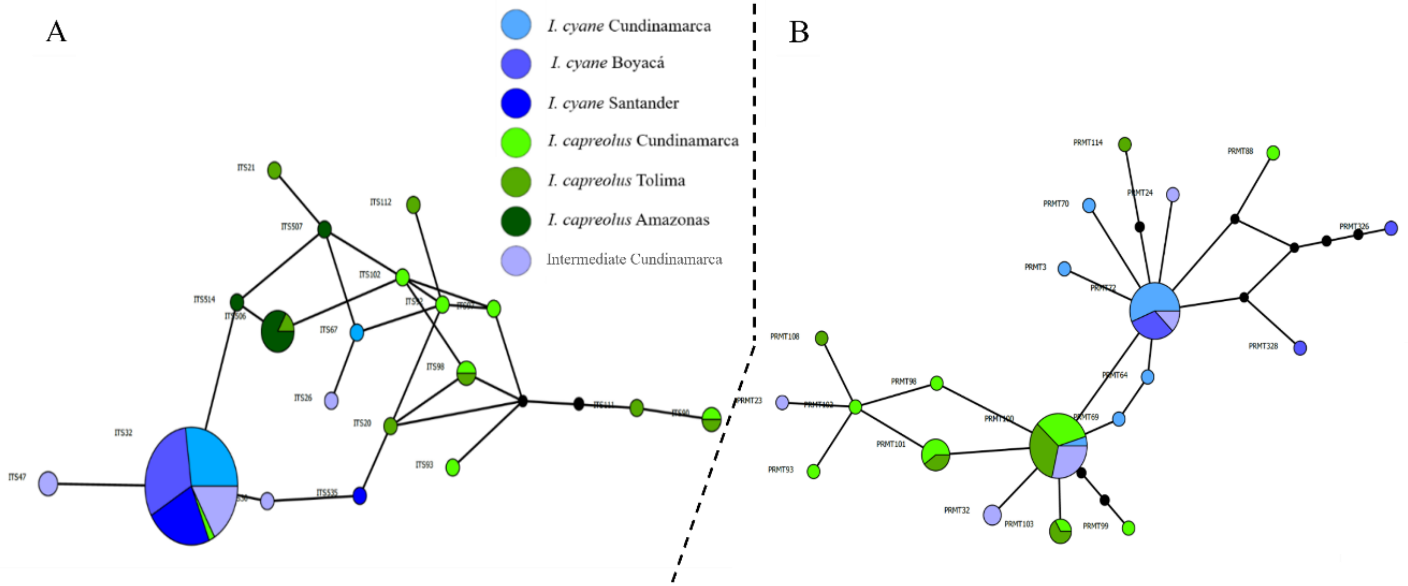

**Figure S6.** Thin-plate spline deformation grids showing the consensus shape for *I. capreolus*, *I. cyane*, and the intermediate population from Anolaima.

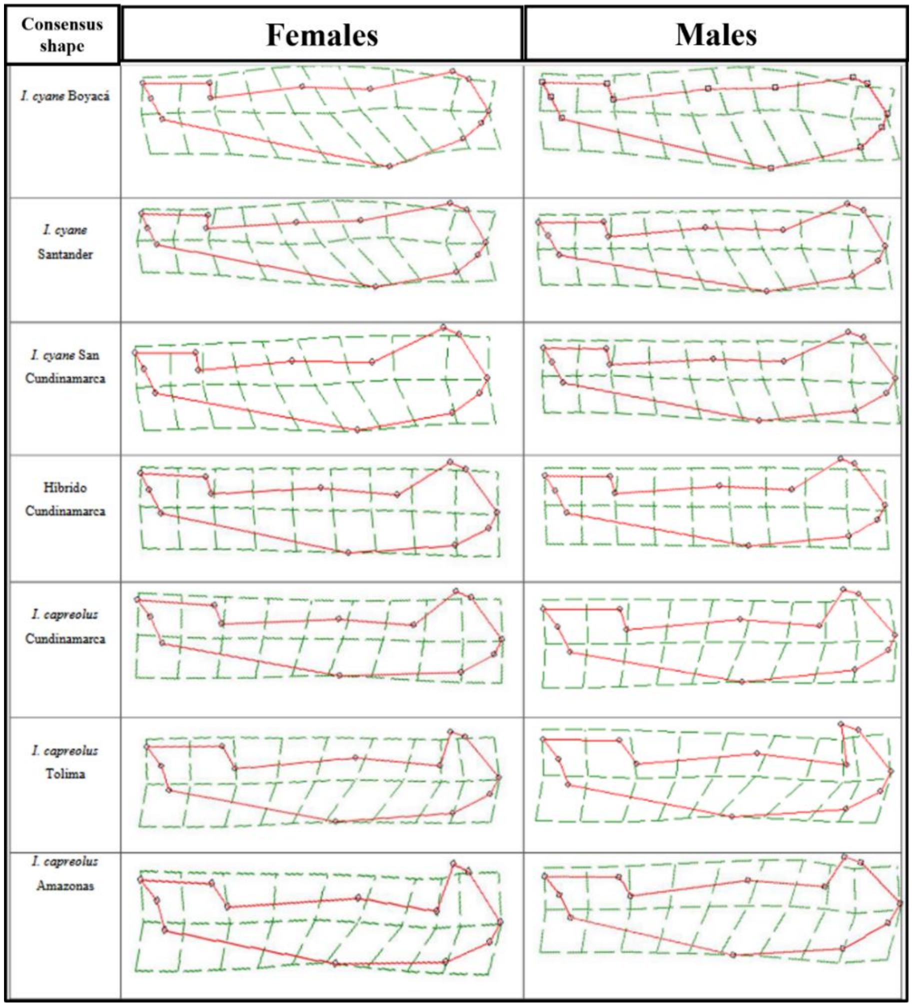

**Figure S7.** Thin-plate spline deformation grids showing the consensus shape of male caudal appendages in *I. capreolus* (left), *I. cyane* (right), and putative hybrids from the Anolaima population (center).

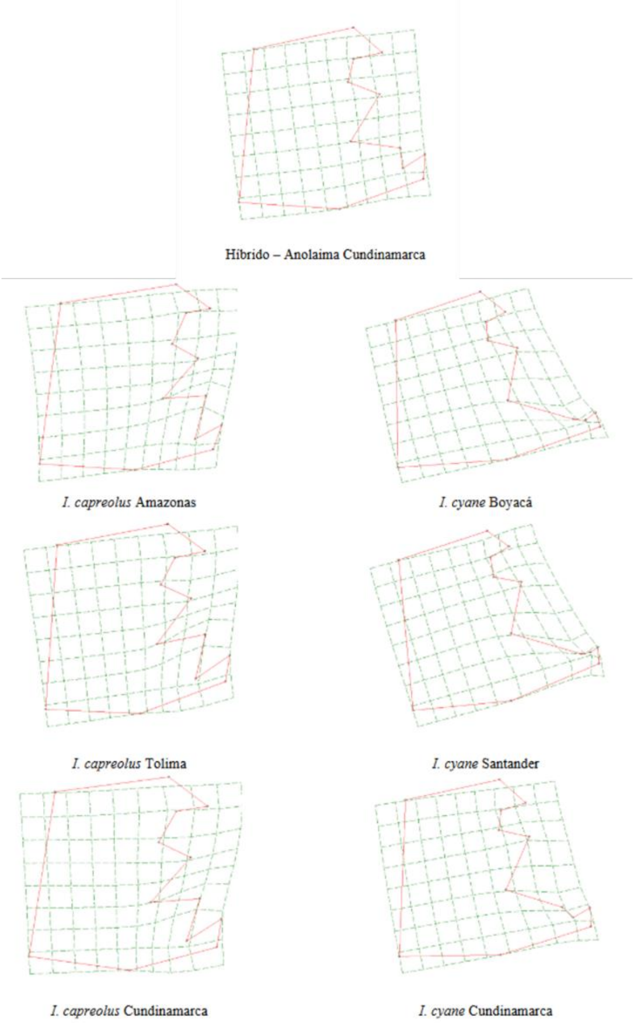
